# Efficacy of Doxycycline as monotherapy or in combination with Levofloxacin in treating late-stage systemic anthrax in non-human primates

**DOI:** 10.1101/2025.07.07.663498

**Authors:** Amir Ben-Shmuel, Assa Sittner, Itai Glinert, Elad Bar-David, Josef Schlomovitz, Haim Levy, Shay Weiss

**Affiliations:** Department of Infectious Diseases, Israel Institute for Biological Research, Ness-Ziona, Israel

**Keywords:** keyword 1, *Bacillus anthracis* 2, Doxycycline 3, Levofloxacin 4, Non-Human Primates 5, Anthrax 6, Treatment 7, Trigger to treat

## Abstract

*Bacillus anthracis* spores pose a major biosecurity threat, in light of their use in several documented bio-terror attacks and the fact that it was weaponized. The absence of specific and accurate treatment instructions in the 2001 anthrax letter attacks and the poor prognosis of patient receiving noneffective treatment, emphasized the need for reliable guidelines for treating exposed(prophylaxis) or symptomatic populations. The recent 2024 CDC guidelines for treating systemic anthrax recommend a combined treatment of a tetracycline (Minocycline or Doxycycline) with Meropenem or a fluoroquinolone. These recommendations differ from the 2001 or 2015 updates, and in the absence of significant clinical data, rely on animal studies. Previously we demonstrated, based on a rabbit model, that these recommendations are efficient in treating various stages of anthrax, ranging from post exposure prophylaxis, through systemic to involvement of the CNS. Herein we confirm our rabbit results by demonstrating the efficacy of Doxycycline as a monotherapy or in combination with Levofloxacin in treating late-stage systemic anthrax in non-human primates (NHP), in a trigger to treat type of experiment. Our experiments show high efficacy of the mono or combined therapy. Altogether, these results support the CDC guidelines by demonstrating the efficacy of this treatment in a second highly relevant NHP model.

## 1. Introduction

*Bacillus anthracis*, a gram-positive spore forming bacterium, is the etiological cause of anthrax, a lethal zoonotic disease that is naturally transmitted to humans by contact with sick animals or contaminated animal products [1-3]. In most cases the spores (the infectious form), penetrate the body through skin lesions, or the digestion or inhalation tracts [1]. The route of infection determines the type of anthrax that develops i.e., local eschar, gastrointestinal and systemic disease. Systemic disease is almost always fatal without treatment and will always develop after inhalational and oral infections [4]. The cutaneous form is the most obvious of the three and will resolve in 80-95% of the cases [5]. A relatively new form of anthrax, deep tissue infection, resulting from the injection of spore-contaminated heroin, causes extended edema and inflammation of the injected limb [6]. Historically, anthrax patients were treated with injections of penicillin G, that were highly effective but of limited applicability in the case of mass exposure event [1]. Following the 2001 anthrax mail attacks and the confusion in treatment of systemic patients, the Centers for Disease Control and Prevention (CDC) recommended treatment was based on Ciprofloxacin or Doxycycline in combination with up to four additional substances [7]. The CDC guideline addressed two major anthrax stages; early-stage post exposure non-symptomatic population (Post exposure prophylaxis - PEP) and symptomatic, most likely systemic patients. The systemic patients’ symptoms could range from flu like to acute condition presenting high fever, plural fluids accumulation and meningitis [8]. For PEP the CDC recommended 60 days of oral treatment with ciprofloxacin or doxycycline. For symptomatic patients the CDC recommended combined treatment based on Ciprofloxacin or Doxycycline together with one or more substances from a list. Eventually the combination of Ciprofloxacin with Clindamycin was the treatment of choice and was successfully used in several anthrax cases since [8].

Since the original publication of guidelines (in 2001), the CDC updated its recommendations twice - in 2015 and 2023 [9,10] . The 2015 recommendations divided the symptomatic patients into two groups based on the physician ability to exclude CNS involvement. The treatment in this case was based on the fluoroquinolone (Ciprofloxacin or Levofloxacin) supplemented with protein synthesis inhibitor (Clindamycin or Linezolid). In the case where meningitis could not be excluded, the CDC recommended combining three antibiotics, a fluroquinolone, a protein synthesis inhibitor (Linezolid as first choice) and a β-lactam (Meropenem). Unlike the 2015 recommendations that were based mainly on data of other CNS diseases, the 2023 update was based on comprehensive analysis of all the available data (published and unpublished) on anthrax treatment in relevant animal models [11]. The recommended treatment is based on a tetracycline (Minocycline or Doxycycline) combined with Meropenem or a fluoroquinolone (Ciprofloxacin or Levofloxacin) [10]. Previously we reported the high efficacy of Doxycycline in treating late stages of systemic anthrax in rabbits as a monotherapy or in combination with a fluoroquinolone or Meropenem [12-15]. In this work we validate the rabbit results in non-human primates (NHP) that were infected via intranasal spray of spores in trigger-to-treat experiments with Doxycycline alone or with Levofloxacin.

## 2. Materials and Methods

This study was carried out by trained personnel, in strict accordance with the recommendations of the Guide for the Care and Use of Laboratory Animals of the National Research Council. All protocols were approved by the IIBR committee on the Ethics of Animal Experiments (Protocol numbers, RM-01-18 and RM-03-19). Animals had free access to food and water, maintained at a controlled environment with constant fresh air supply and ambient room temperature of 24° C. Enrichment was supplied in the form of specialized toys. For each experiment animals were picked randomly, sedated using 100mg ketamine and 10mg xylazine and transferred from the general animal facility to the BSL3 facility for acclimation. As the animal transfer between the general animal facility and the BSL3 is unidirectional, to avoid euthanizing naïve animals, the negative control animals were maintained in the general animal facility and were monitored daily by the facility staff.

Animals (female Rhesus macaque) were anesthetized using 100mg ketamine and 10mg xylazine and infected by spraying 1×10^6^ CFU (10xLD_50_) *B. anthracis* Vollum spore suspension [13] into their nasal cavity with a MicroSprayer® Aerosolizer for Rats — Model IA-1B-R (PennCentury™). Blood samples (3 ml) were drawn from a limb vein every 4h starting 24 hours post inoculation, and serum protective antigen (PA) concentration was determined using a home-made rapid ELISA [14] as a correlate for bacteremia level (Table 1). When bacteremia was estimated as >10^4^ CFU/ml blood was drawn and plated to determine the exact bacteremia (CFU/ml) and treatment was initiated. As we previously demonstrated in rabbit and guinea pig experiments, this level of bacteremia represents progressed systemic disease at a stage that effective antibiotic treatment will be successful in >95% of the cases and dropped dramatically when bacteremia is higher than 1×10^6^ CFU/ml.

**Table 1.** Serum PA and bacteremia at treatment initiation.

| Bacteremia (CFU/ml) | Serum PA ($\mu\text{g/ml}$) |
| --- | --- |
| $3 \times 10^8$ | $>50$ |
| $1.2-4 \times 10^7$ | 20-30 |
| $1-2 \times 10^6$ | 3-5 |
| $3-5 \times 10^5$ | 0.3-0.5 |
| $1-2 \times 10^4$ | 0.05 |

Animals were treated SC with Doxycycline (15 mg/kg) as a monotherapy or combined with Levofloxacin (20 mg/kg) twice a day for three days, followed by doxycycline alone, SC twice a day for four days. Animals were monitored twice a day for 14 days post inoculation for behavioral signs and body temperature using a subcutaneous RFID chip in the first 7-8 days. Animals unable or unwilling to drink were injected with 20-100ml of saline or dextrose isotonic solution SC. Animals were euthanized immediately by a 120 mg/kg sodium pentobarbitone injection when one of the following symptoms was detected: severe respiratory distress or the loss of righting reflex. Since in most cases anthrax symptoms are detectable only in close proximity to death, both animals succumbed to the disease. To reduce the use of NHP these two animals, one from each treatment group, serves as a positive control.

The use of *B. anthracis* was performed according to the Israeli select agent law. The experiments were regulated and monitored by the head of the biosafety unit and by the institutional regulatory committee at the IIBR. All experiments were conducted in a BSL3 laboratory.

## 3. Results

The Doxycycline monotherapy group consisted of six animals. The animals were sedated and infected by spraying 1×10^6^ CFU (10xLD_50_) *B. anthracis* Vollum spore suspension [13] into their nasal cavity with a MicroSprayer® Aerosolizer. 24 h post infection, blood samples were obtained, and PA serum concentration was estimated using a previously reported rapid ELISA [16]. In this specific experiment all the animals were positive for serum PA at levels that indicate bacteremia at the desired levels (Table 1). At this point we decided to start treating all the animals in this group, though according to our prediction we estimated would have bacteremia lower than the target of 1×10^4^ CFU/ml, since according to our experience at these bacteremia levels, deterioration could be fast. The precise bacteremia of each of the animal at treatment initiation was determined by plating diluted blood samples. In perspective we treated animals with bacteremia between 200 to 3×10^8^ CFU/ml (Fig 1A). Animals were treated twice a day SC with 15 mg/kg of Doxycycline. Animal pose and general behavior was observed from distance using in room camera, and body temperature was recorded daily. Similar to the efficacy reported in the rabbit model [14], Doxycycline monotherapy was highly effective, protecting all the animals with bacteremia of up to 10^7^ CFU/ml, but failing to protect the animal with bacteremia of 3×10^8^ CFU/ml. Body temperature in all surviving animals (Fig 2A) showed minimal differences throughout the treatment period. In contrast, there was a rapid decrease in temperature of the animal that succumbed to the infection. This animal died within 36h from treatment initiation, it’s body temperature dropping to 32°C within 24h and succumbing within the following 12 h, receiving two rounds of antibiotic treatment in the duration. Gross necropsy revealed high accumulation of plural fluids and high intracranial tension that included hemorrhage, estimated by the excessive amount of bloody CSF discharged during brain necropsy. Both blood and brain were sterile, which is similar to what we had previously demonstrated for clindamycin in the rabbit model [16]. We cannot exclude the presence of bacteria in the organs, however the fact that the blood and CSF were sterile indicate that the death was not due to treatment failure.

**Figure 1.**
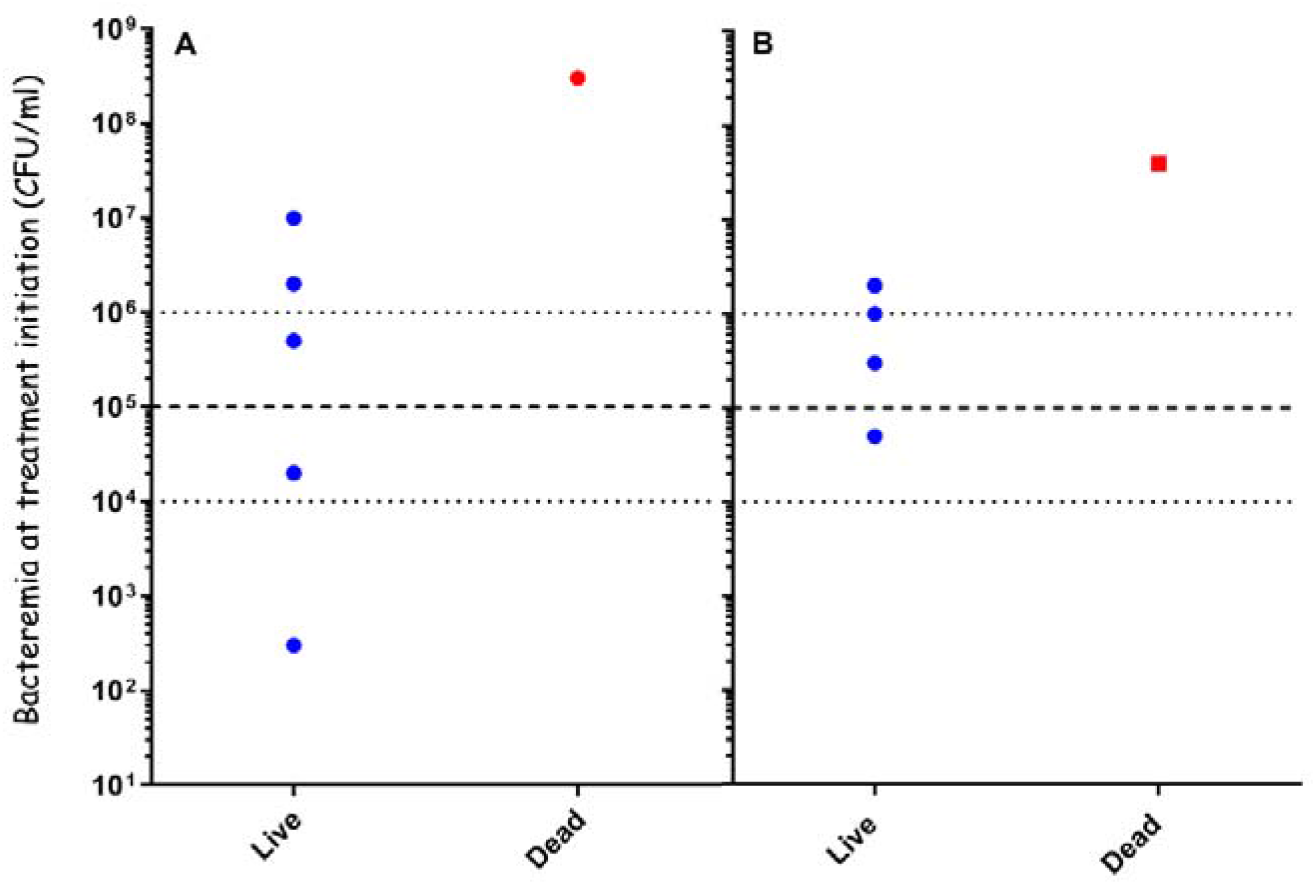
Outcome of inhalation anthrax treatment with Doxycycline alone or in combination with Levofloxacin. Animals were infected intranasally with Vollum spores using a PennCentury™ device. Disease progression was monitored by determining serum PA concentration which served as the trigger to treat. Bacteremia (CFU/ml) at treatment initiation and outcome the treatment (Live or Dead) is indicated for A – Doxycycline and B – Doxycycline + Levofloxacin.

**Figure 2.**
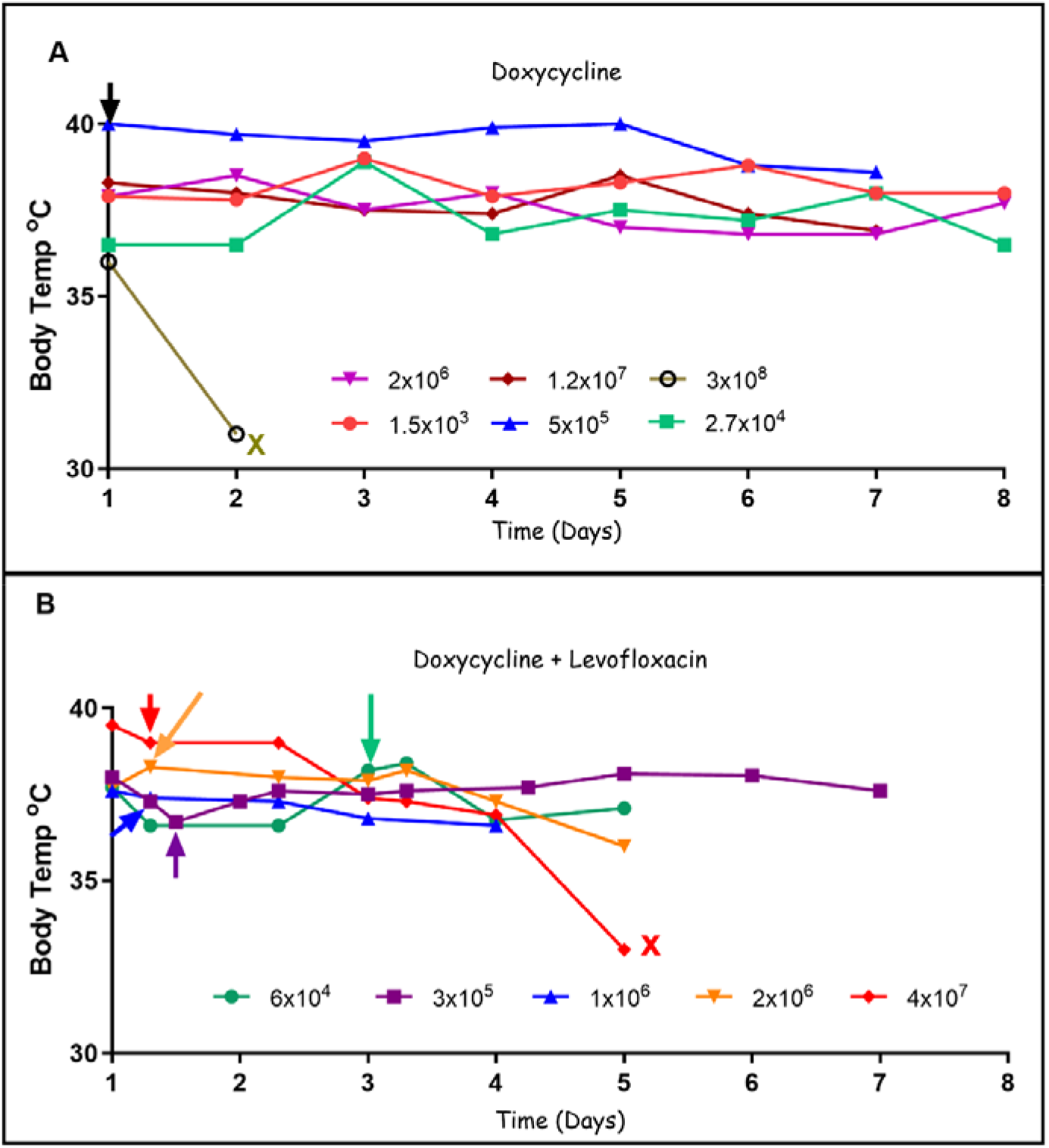
Body temperature of individual animals during the treatment period. A subcutaneous chip was used to record body temperature throughout the treatment period. A – Doxycycline and B – Doxycycline + Levofloxacin. Arrows indicate treatment initiation and X indicated that the specific animal succumbed to the disease. The Doxycycline treatment was initiated at 24 h (black arrow) post infection, while in the Doxycycline + Levofloxacin experiment the treatment onset variation was high and the arrows are color coded according to the specific animal treated. The Bacteremia at treatment initiation is indicated in the legend.

Since in the case of systemic disease the combination of at least two antibiotics is recommended [10], we tested the effect of adding Levofloxacin to the treatment. For this experiment we expose five animals were exposed to *B. anthracis* Vollum spores as described for the Doxycycline mono-treatment. Serum PA was estimated every 4h starting 24h post infection. Unlike the first experiment, the disease progressed differently in each of the animals, resulting in treatment initiation at different time points, as shown by collared arrows in figure 2B. This treatment group consist of five animals with bacteremia of 5×10^4^ to 4×10^7^ CFU/ml. Animals were treated twice a day SC with 15 mg/kg of Doxycycline and 20 mg/kg of Levofloxacin for three days followed by a four days treatment with Doxycycline as mon-therapy. Animals ware monitored as previous This combined treatment was highly effective, protecting animals with a bacteremia of 5×10^4^ to 2×10^6^ CFU/ml (Fig 1B).

This treatment did not protect a single animal with 4×10^7^ CFU/ml. Body temperature monitoring (Fig 2B) showed again that in surviving animals no major changes were observed, while there was a slow decrease in temperature during the four days from treatment initiation to death in the animal that succumbed. Once this was detected, a warming infrared lamp was added, and 200 ml of saline were administrated SC twice a day. This limited supportive treatment, on top of the therapeutic protocol, did not prevent the animal’s death. Perhaps, intensive care level of treatment might have been more effective in preventing animal’s death. Gross necropsy did not revile any obvious damage to internal organs and there was no bacterial growth in blood or brain samples.

## 4. Discussion

Anthrax is rare lethal zoonotic disease that depends completely on the animal role to develop and validate new medical countermeasures [3]. The recent CDC recommends a combined treatment of tetracycline (Minocycline or Doxycycline) with Meropenem or fluoroquinolone (Levofloxacin) for the treatment of a systemic anthrax [10]. Since previously, tetracycline was limited to PEP [9], most of the published data refers to the efficacy of this drug in treating animals in 24h post exposure. At this time point most of the animal do not develop any sign of disease and are probably not bacteremic. Since animal do not show any disease signs the only reliable marker is blood bacteremia which increases with the progression of the disease, when close to death can reach 10^8^-10^9^ CFU/ml [17]. Therefore, when testing treatment efficacy, a trigger to treat type of experiments are used. Since live counts are time consuming and are relevant only in retrospect, in most experiments, plasma toxins, mainly PA, are used as trigger to treat marker [14,18-20]. Since the correlation of bacteremia to PA is not perfect, in most of these experiments, treatment is initiated immediately once PA is detected, and is usually based on large cohort of animals to achieve statistical significance[21-23]. Therefore, the accepted model for such experiments is the rabbit, however, NHP are still considered the relevant animal model when translating scientific data to human recommendations [24].

The efficacy of anthrax treatment is in correlation to the time from infection, where the efficacy decreases as the disease progresses. Treating exposed individuals (non symptomatic) is most effective. In 2001 anthrax letter attack thousands of suspected and confirmed people, received ciprofloxacin or doxycycline prophylaxis, none of them develop anthrax symptoms [25]. Modeling treatment of symptomatic individuals is more complicated since it includes variation of patients exhibiting variety of signs ranging from flu like to acute phase, which includes central nerves system (CNS) involvement [8]. The aim of this work was to evaluate the efficacy of the recent CDC guidelines in treating systemic anthrax patients, with or the absence of indications of CNS involvement [10]. For this purpose, we can use two types of experiments: direct CNS infection model [26] and a trigger to treat type of experiment following airway spore infection. In the trigger to treat type of experiments we aim to bacteremia ranging from 0 to 10^8^ CFU/ml, one that will allow us to distinguish between different stages of the systemic disease. Based on our pervious finding indicating that from a bacteremia of 10^5^ CFU/ml bacteria could be detected in the brain tissue [17], we defined bacteremia of 10-10^4^ CFU/ml as flu like and 10^4^-10^6^ CFU/ml as acute stages of the disease. In our rabbit experiments we defined the treatment as efficient when all the animals with bacteremia of up to 10^4^ at treatment initiation were saved [15,16].

Previously we demonstrated that Doxycycline, both as monotherapy or combined with Levofloxacin, is effective in treating late-stage systemic disease or following direct CNS infection in rabbits [12,14]. We also demonstrated that this treatment is effective in NHP following direct CNS infection [12]. Since the CDC guidelines are referring to the treatment of patients with progressed systemic anthrax, the NHP experiment had to simulate this condition. The disease that develops following CSF injection is mainly a CNS disease [26] while the systemic organ damage is secondary and results from transfer from the CNS to the bloodstream (the opposite direction of the natural disease). This model is great in testing the ability of antibiotics to cross the BBB and be effective in resolving the infection [26,27]. In this experiment we started with a pulmonary infection of the animal and monitored the PA level carefully in an attempt to get maximal bacteremia variation using a limiting number of animals, since we are dealing with NHP. All the animals presented with bacteremia at treatment initiation, indicating that they were all successfully infected and developed a disease that was lethal in the absence of effective revetment. The fact the one animal from each treatment group succumbed to the infection, while being treated with antibiotics, supports this statement and can be considered as a positive control of sorts. Herein we tested the efficacy of Doxycycline as a mono treatment or in combination with Levofloxacin as a treatment of late-stage systemic disease following airway spore infection in NHP. The two treatments were highly effective, protecting animals with bacteremia as high as 10^6^-10^7^ CFU/ml. The fact that there is one animal in each bacteremia level restricts our ability of making statistical claims regarding the differences between the two groups. However the fact that five animals in the Doxycycline group, with bacteremia of up to 1×10^7^ CFU/ml and four animals from the group treated with Doxycycline and Levofloxacin with bacteremia of up to 2×10^6^ CFU/ml, survived, indicates high treatment efficacy. Our treatment failed to protect one animal from each group. Doxycycline failed to protect an animal that had a very high bacteremia level of 3×10^8^ CFU/ml. This bacteremia level is typical to moribund animals, and is impossible to treat by antibiotic treatment. The finding of large quantities of plural fluid and high intracranial pressure due to CSF accumulation, similar to that reported for human anthrax patients, supports this assumption. The fact that the blood and CSF were sterile after two doses of antibiotics, points to the potency of Doxycycline and not that the animal is actually sterile. The animal that died during the Doxycycline and Levofloxacin treatment had a high bacteremia of 4×10^7^ CFU/ml at treatment initiation, and actually improved during the first three days of treatment. However, the overall condition of the animal deteriorated and our supportive treatment of fluids and heating lamp was insufficient and the animal died without any obvious necropsy finding. Since this animal received five days of antibiotic treatment, we can assume that the animal was sterile. In these two cases we can assume that intensive care treatment would prolong the survival of these animals, especially the one who died on day six. Overall, These findings support our previous rabbit results and the recent CDC recommendations for treating this stage of the disease.

## Author Contributions

Conceptualization, H.L., S.W. and A.B.S.; methodology, E.B.D, J.S. and I.G.; validation, S.W., A.S., and H.L.; formal analysis, H.L. and S.W.; investigation, A.B.S., A.S., I.G., E.B.D., J.S., H.L. and S.W.; writing—original draft preparation, H.L., S.W., A.B.S. and I.G; writing—review and editing, A.S., E.B.D. and J.S..; visualization, H.L. and S.W..; supervision, H.L..; project administration, H.L. All authors have read and agreed to the published version of the manuscript.

## Institutional Review Board Statement

This study was carried out by trained personnel, in strict accordance with the recommendations of the Guide for the Care and Use of Laboratory Animals of the National Research Council. All protocols were approved by the IIBR committee on the Ethics of Animal Experiments (Protocol numbers, RM-01-18 and RM-03-19).

## Informed Consent Statement

The study did not involve humans

## Data Availability Statement

All the relevant data is presented in the manuscript Conflicts of Interest: The authors declare no conflict of interest

